# Strongly coupled transmembrane mechanisms control MCU-mediated mitochondrial Ca^2+^ uptake

**DOI:** 10.1101/2020.04.07.029637

**Authors:** Horia Vais, Riley Payne, Carmen Li, J. Kevin Foskett

**Affiliations:** Departments of Physiology, University of Pennsylvania, Philadelphia, PA, USA; Departments of Cell and Developmental Biology, Perelman School of Medicine, University of Pennsylvania, Philadelphia, PA, USA

## Abstract

Ca^2+^ uptake by mitochondria regulates bioenergetics, apoptosis, and Ca^2+^ signaling. The primary pathway for mitochondrial Ca^2+^ uptake is the mitochondrial calcium uniporter (MCU), a Ca^2+^-selective ion channel in the inner mitochondrial membrane. MCU-mediated Ca^2+^ uptake is driven by the sizable inner-membrane potential generated by the electron-transport chain. Despite the large thermodynamic driving force, mitochondrial Ca^2+^ uptake is tightly regulated to maintain low matrix [Ca^2+^] and prevent opening of the permeability transition pore and cell death, while meeting dynamic cellular energy demands. How this is accomplished is controversial. Here we define a regulatory mechanism of MCU-channel activity in which cytoplasmic Ca^2+^ regulation of intermembrane space-localized MICU1/2 is controlled by strongly-coupled Ca^2+^-regulatory mechanisms localized across the membrane in the mitochondrial matrix. Ca^2+^ that permeates through the channel pore regulates Ca^2+^ affinities of coupled inhibitory and activating sensors in the matrix. Ca^2+^ binding to the inhibitory sensor within the MCU amino-terminus closes the channel despite Ca^2+^ binding to MICU1/2. Conversely, disruption of the interaction of MICU1/2 with the MCU complex abolishes matrix Ca^2+^ regulation of channel activity. Our results demonstrate how Ca^2+^ influx into mitochondria is tuned by coupled Ca^2+^-regulatory mechanisms on both sides of the inner mitochondrial membrane.

---

The mitochondrial Ca^2+^ uniporter (MCU) is a Ca^2+^-selective ion channel in the inner mitochondrial membrane (1) that represents the major mechanism by which mitochondria take up Ca^2+^ into the matrix to regulate bioenergetics, cell death pathways and cytoplasmic Ca^2+^ signaling. MCU-mediated Ca^2+^ uptake is driven by the large (−150 – -180 mV) inner-membrane voltage (ΔΨ_m_) generated by proton pumping into the intermembrane space (IMS) by the electron-transport chain during oxidative phosphorylation. Despite the considerable driving force for Ca^2+^ entry into the mitochondrial matrix, mitochondrial matrix free [Ca^2+^] ([Ca^2+^]_m_) at rest is maintained at ∼100 nM, similar to the resting cytoplasmic free [Ca^2+^] ([Ca^2+^]_i_). Whereas transient elevations of [Ca^2+^]_m_ promote enhanced ATP production to support cellular energy demands, failure to maintain low matrix [Ca^2+^] may trigger ΔΨ_m_ depolarization, and opening of the permeability transition pore (PTP) that can result in cell death. Thus, mitochondrial Ca^2+^ uptake must be tightly controlled, but how mitochondrial Ca^2+^ uptake through MCU is regulated is controversial (2, 3). The MCU channel in metazoans exists as a protein complex comprised of MCU as the tetrameric channel pore-forming subunit (4, 5), EMRE, a single-pass transmembrane protein that is essential for channel activity (6-8), and IMS-localized (but see (9)) Ca^2+^-binding EF hand domain-containing MICU1/2 heterodimers (10-13) that associate with the channel complex by interactions with the carboxyl-terminus of EMRE (14) and with MCU (15, 16). In a current model of MCU regulation (but see (2)), MICU1/2 promotes cooperative channel activation in response to Ca^2+^ binding by their EF hands, whereas apo-MICU1/2 imposes a threshold [Ca^2+^]_i_ below which MCU channel activity is inhibited, so-called channel gatekeeping (13, 17-21).

Two fundamental observations suggest that this model of MCU regulation is inadequate. First, it cannot account for the observation that the channel can be open with the MICU1/2 EF-hands in their apo state under conditions in which [Ca^2+^]_i_ is reduced to low levels that permit Na^+^ permeation (1, 22, 23). Second, by mitoplast electrophysiology of the inner mitochondrial membrane (IMM) with pipette solutions containing defined free-[Ca^2+^], it was observed that matrix [Ca^2+^] also regulates MCU-channel gating, with strong inhibition at normal resting [Ca^2+^]_m_ and maximal channel inhibition ∼80% at ∼400 nM (23). In this condition, the channel was strongly inhibited despite MICU1/2 being fully Ca^2+^-liganded by Ca^2+^ present in the bathing solution, an observation that also cannot be accounted for by the current model of MCU regulation.

Here we set out to understand the matrix [Ca^2+^] regulation of MCU-channel activity. By varying mitochondrial Ca^2+^ buffering capacity and [Ca^2+^]_m_ in mitoplast patch-clamp electrophysiology experiments, we determined that Ca^2+^ that permeates through the channel pore regulates Ca^2+^ affinities of coupled inhibitory and activating sensors in the matrix. Ca^2+^ binding to the inhibitory sensor that we have localized within the MCU amino-terminus closes the channel despite Ca^2+^ binding to MICU1/2. Disruption of the this inhibitory Ca^2+^-binding site promotes mitochondrial Ca^2+^ overload and PTP opening in intact mitochondria in response to Ca^2+^ additions. Conversely, disruption of the interaction of MICU1/2 with the MCU complex abolishes matrix Ca^2+^ regulation of channel activity. A model of coupled transmembrane Ca^2+^-regulatory mechanisms can account for these data, as well as for the observation that the channel can be open with the MICU1/2 EF-hands in their apo state under conditions in which Na^+^ is the permeant ion (1, 22, 23).

## Results

### Matrix [Ca^2+^] Regulation of MCU Ca^2+^ Channel Activity Coupled to MICU1/2 Regulation

We previously determined that matrix [Ca^2+^] regulated MCU-channel open probability with a biphasic concentration dependence, with strong inhibition at normal resting [Ca^2+^]_m_ (∼100 nM) and maximal channel inhibition ∼80% at ∼400 nM (23). We first confirmed this matrix [Ca^2+^] regulation by recording MCU Ca^2+^ currents in mitoplasts isolated from HEK293 cells, with the patch-clamp pipette solution that filled the mitochondrial matrix containing defined [Ca^2+^] buffered with 1.5 mM EGTA. Ca^2+^ currents were large with pipette solutions containing 1.5 mM EGTA with either no added Ca^2+^ or with free [Ca^2+^] set to 1 µM, whereas they were inhibited by ∼80% at 400 nM Ca^2+^, in excellent agreement with the previous observations (Fig. 1*A, E*). To explore the relationship between matrix [Ca^2+^] regulation and MICU1/2-mediated cytoplasmic [Ca^2+^] regulation of MCU activity, we genetically knocked down MICU1 or MICU2, or inhibited their association with the channel complex by mutating the EMRE carboxyl-terminus (14). We previously demonstrated that knockdown of either MICU1 or MICU2 or expression of mutant EMRE have no effects on MCU expression (18). In agreement, Ca^2+^-current densities measured with 0- or 1 µM Ca^2+^ in the pipette solution were comparable in MICU knockout and mutant EMRE-expressing cells to those in wild-type cells (Fig. 1*C* and *D*). However, each of these genetic interventions that abolish [Ca^2+^]_i_ regulation of MCU channel activity also abolished matrix [Ca^2+^] regulation (Fig. 1*C*; see also (23)). Thus, channel regulation by matrix [Ca^2+^] and by MICU1/2 on the opposite side of the IMM are strongly coupled. Accordingly, in the simplest model to account for MCU channel regulation, the channel complex possesses two matrix-localized Ca^2+^-sensing sites, inhibitory (S_i_) and activating (S_a_), whose Ca^2+^ occupancies control the functionality of MICU1/2 on the other side of the membrane to regulate channel gating.

**Fig. 1.**
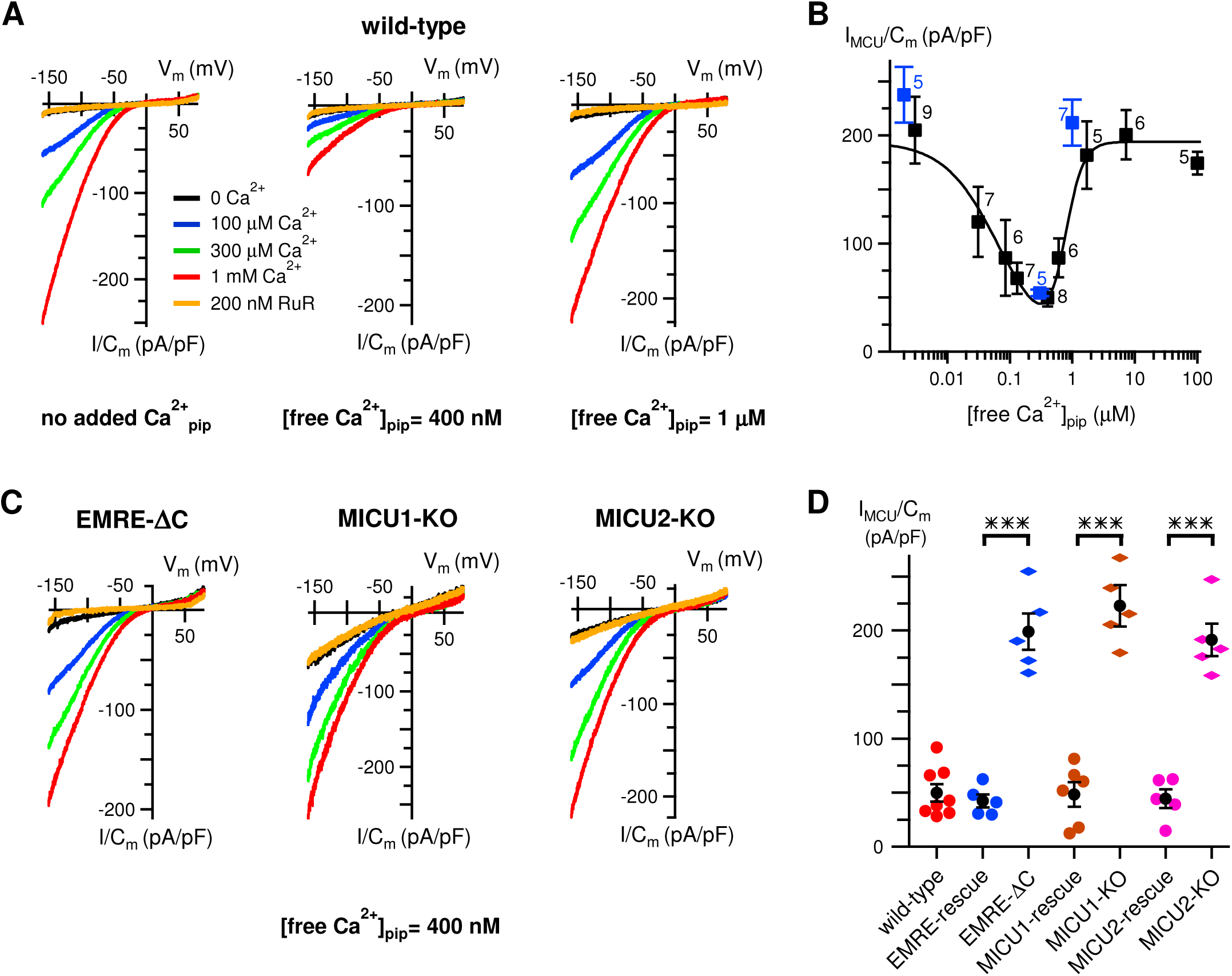
Matrix [Ca^2+^] regulation of MCU requires the MICU1/2 heterodimer. (*A*) Representative families of MCU Ca^2+^ currents in response to voltage ramps in wild-type mitoplasts, with pipette free [Ca^2+^] in 1.5 mM EGTA indicated under each panel. Colors denoting bath conditions are used throughout. (*B*) Biphasic matrix [Ca^2+^] regulation of MCU Ca^2+^-current density in wild-type mitoplasts at -160 mV with 1 mM bath [Ca^2+^]. Data from the current study shown in blue. Data from ref. (23) shown in light gray for reference. Peak inhibition is observed with matrix free [Ca^2+^] = 400 nM. Numbers indicate number of mitoplast recordings. Bars denote standard error of the mean (SEM). (*C*) Representative MCU Ca^2+^ currents recorded with pipette free [Ca^2+^] = 400 nM in mitoplasts from cells lacking functional MICU1/2 regulation due to expression in EMRE knockout (KO) cells of EMRE with its acidic carboxyl-terminus deleted (EMRE-ΔC) (left), or in MICU1-(middle) or MICU2-(right) KO cells. (*D*) Summary of MCU Ca^2+^-current densities recorded with matrix free [Ca^2+^] = 400 nM in mitoplasts from wild-type cells, and EMRE-ΔC, MICU1-(middle) or MICU2-(right) KO cells, and their respective rescue cells. Loss of normal inhibition observed with matrix free [Ca^2+^] = 400 nM in cells lacking MICU1/2 regulation is rescued by expression of wild-type EMRE, MICU1 and MICU2. Bars: SEM, \*\*\**P* < 0.00003. See also ref. (23).

### Matrix Ca^2+^ Buffering Regulates Matrix Ca^2+^ Regulation of MCU Ca^2+^ Channel Activity

Whereas it was previously inferred that the EMRE carboxyl-terminus could contribute to matrix Ca^2+^ sensing (23), its location in the IMS (7, 14) precludes a direct role. To identify matrix Ca^2+^-regulatory sites, we first considered that plasma membrane Ca^2+^ channels can be regulated by [Ca^2+^]_i_ in the nanodomain created by Ca^2+^ flux through the channel (24). Inhibition of such regulation by the fast Ca^2+^ buffer BAPTA but not by a slow buffer such as EGTA is considered evidence for permeant Ca^2+^ ion-flux regulation of channel activity (24, 25). To determine whether Ca^2+^ flux through MCU contributes to matrix [Ca^2+^] regulation of its activity, we replaced 1.5 mM EGTA with the same concentration of BAPTA. Whereas we predicted that faster buffering might either eliminate matrix [Ca^2+^] regulation or shift the inhibitory [Ca^2+^]_m_ to higher concentrations, BAPTA unexpectedly shifted the entire biphasic matrix [Ca^2+^] dependence to 10-fold lower concentrations (Fig. 2*A*). Conversely, reducing [BAPTA] from 1.5 mM to 0.5 mM shifted the biphasic dependence in the opposite direction (Fig. 2*B*). Incremental increases of [BAPTA] in the pipette solution shifted the biphasic matrix [Ca^2+^] dependence of MCU activity to lower concentrations, such that at 5 mM BAPTA only the maximal inhibition and recovery from inhibition could be observed (Fig. 2*A* and *B*). A linear relationship in a semi-log plot of [BAPTA] and the inhibitory [Ca^2+^]_m_ (Fig. 2*C*) suggests that matrix Ca^2+^ buffering regulates a single dominant process that controls matrix [Ca^2+^] regulation of channel activity. Physiologically, mitochondrial matrix Ca^2+^ buffering capacity is provided by phosphate (P_i_) (26). Matrix [P_i_] also modulated [Ca^2+^]_m_ regulation of MCU-channel activity (*SI Appendix Fig. S1*).

**Fig. 2.**
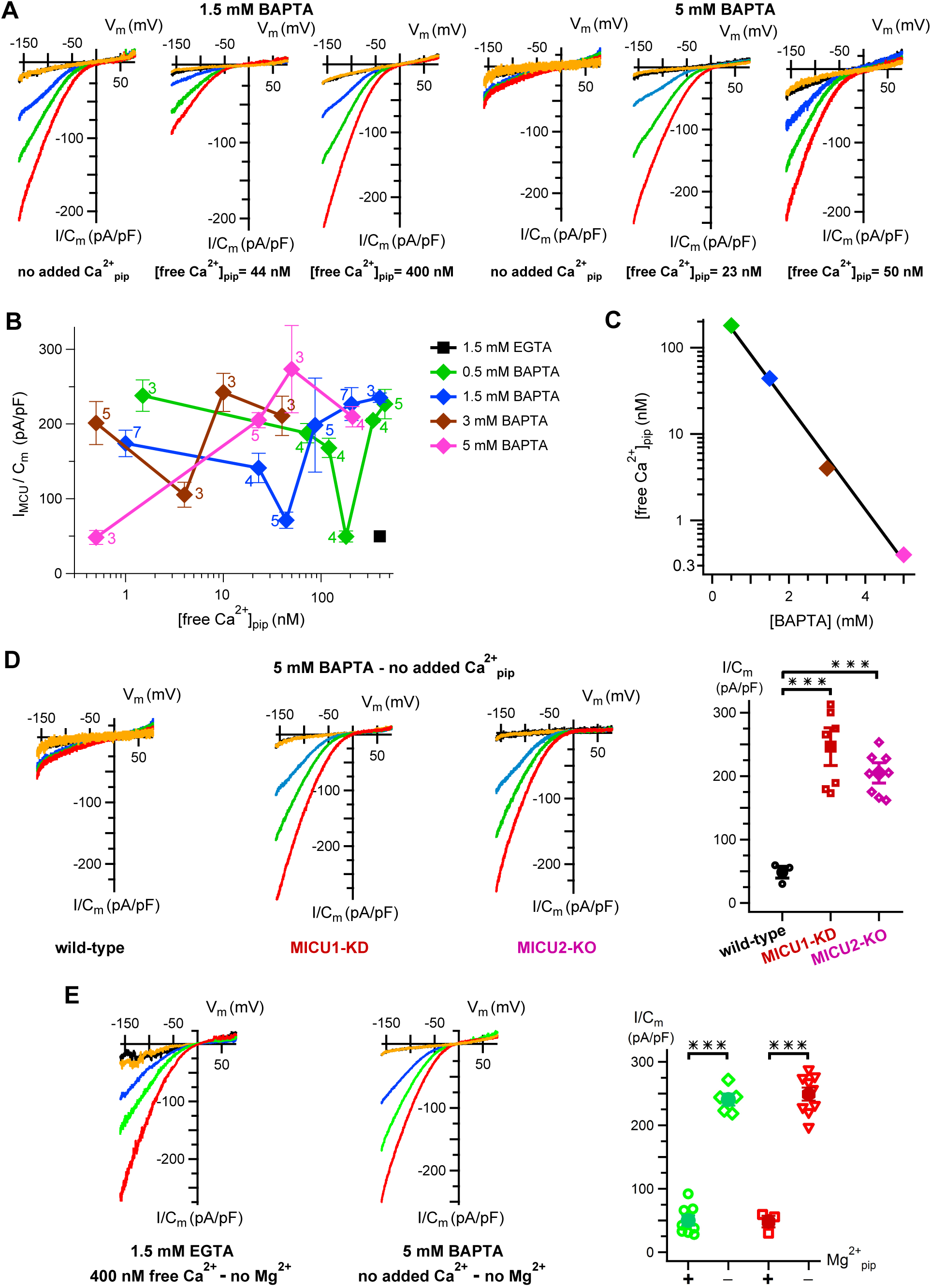
[Ca^2+^]_m_ regulation of MCU is sensitive to matrix buffering capacity. (*A*) Representative MCU Ca^2+^ currents with different pipette free [Ca^2+^] (shown below each set of traces) buffered with either 1.5 mM BAPTA (left group of three panels) or 5 mM BAPTA (right group of three panels). Normal inhibition with matrix free [Ca^2+^] = 400 nM buffered with 1.5 mM EGTA is shifted to 10-fold lower concentration in presence of 1.5 mM BAPTA (center panel in the left group). In 5 mM BAPTA, inhibition is shifted further to lower matrix free [Ca^2+^] (left panel in the right group). (*B*) MCU Ca^2+^-current density over range of pipette free [Ca^2+^] and buffering capacity. Black symbol marks the minimum reported in Fig 1*A*. Numbers indicate numbers of mitoplasts recorded. Biphasic dependence of Ca^2+^-current density on matrix free [Ca^2+^] is shifted to lower concentrations with stronger buffering. (*C*) Relationship between the maximally-inhibited MCU Ca^2+^-current density recorded (from (*B*)) as a function of pipette [BAPTA]. (*D*) Matrix [Ca^2+^] inhibition of MCU Ca^2+^-channel activity observed with pipettes containing 5 mM BAPTA and no added Ca^2+^ (left; see also panel B) requires MICU1 (middle) and MICU2 (right; representative families of current recordings). Right: summary of Ca^2+^-current densities in mitoplasts recorded at -160 mV with 1 mM bath [Ca^2+^] with each point representing a single mitoplast. Bars: standard error of the mean (SEM), ***, *P* = 0.0007 (MICU1) and *P* = 0.00009 (MICU2). (*E*) Inhibition of MCU activity observed with matrix Ca^2+^ = 400 nM buffered with 1.5 mM EGTA (as in Fig. 1*A* and *B*) (left) or with the matrix solution containing 5 mM BAPTA with no added Ca^2+^ (as in panels *B* and *D*) (middle), is abolished in the absence of matrix Mg^2+^. Bars: SEM, ***, *P* < 0.0000009.

Sensitivity to matrix BAPTA and P_i_ of matrix [Ca^2+^] regulation of MCU activity suggests that Ca^2+^ flux through the channel regulates its activity, implying the existence of what we here term a low-affinity Ca^2+^-flux sensor S_f_. Whereas it was expected that BAPTA would de-sensitize the channel to inhibition by matrix [Ca^2+^], the opposite was observed. Notably, the biphasic nature of matrix [Ca^2+^] regulation remained intact under different buffer regimes (Fig. 2*B*). This implies that the inhibitory S_i_ and activating S_a_ Ca^2+^ sensors are coupled and jointly regulated by the Ca^2+^-flux sensor S_f_. The simplest model suggests that Ca^2+^ occupancy of S_f_ tunes the Ca^2+^ affinities of S_i_ and S_a_. When S_f_ is occupied, for example when matrix Ca^2+^ buffering is low, the apparent Ca^2+^ affinities of S_i_ and S_a_ are low, resulting in the biphasic dependence shifted to higher matrix [Ca^2+^]. When S_f_ is unoccupied, for example when the matrix is buffered with 5 mM BAPTA, the apparent Ca^2+^ affinities of S_i_ and S_a_ are high, resulting in the biphasic dependence shifted to very low [Ca^2+^]_m_.

Whereas this model can account for most of the results, strong channel inhibition observed in 5 mM BAPTA at very low [Ca^2+^]_m_ (Fig. 2*A* and *B*) was not predicted, since all matrix Ca^2+^-regulatory sites as well as the flux sensor should be unoccupied, and MICU1/2 was fully Ca^2+^-liganded under our recording conditions, which together would be expected to activate the channel. Matrix [Ca^2+^] inhibition of channel activity observed under these conditions was fully abolished by genetic deletion of MICU1 or MICU2 (Fig. 2*D*), indicating that uncoupling of matrix [Ca^2+^]- and MICU1/2-regulation could not account for channel inhibition observed in this high-[BAPTA]/low-[Ca^2+^]_m_ condition. The pipette solutions in all of our Ca^2+^-current measurements contained 2 mM MgCl_2_, used historically to increase mitoplast electrical stability (22). We therefore considered that Mg^2+^ binding to the inhibitory Ca^2+^ sensor might underlie channel inhibition observed in high-[BAPTA]/low-[Ca^2+^]_m_. Remarkably, not only did removal of Mg^2+^ from the pipette solution completely abolish MCU-channel inhibition observed in 5 mM BAPTA, it also eliminated MCU-channel inhibition normally observed in 400 nM Ca^2+^ /1.5 mM EGTA (Fig. 2*E*).

### Identification of the Matrix Ca^2+^ Inhibition Sensor

Previously, a structure of the isolated amino-terminal domain of human MCU, located in the mitochondrial matrix in the full-length channel (27), revealed an acidic patch with a bound Mg^2+^ that was postulated to be a low-affinity divalent cation-binding site (28). We speculated that this site might be the structural basis for matrix [Ca^2+^] inhibition in full-length MCU. To test this, we individually mutated two key aspartic-acid residues important for divalent-cation binding (28) – D131 and D147 – to alanines (D131A, D147A) and recorded Ca^2+^ currents through the mutant MCU channels following stable expression in MCU-knockout HEK cells over-expressing MICU1 to ensure proper regulation of the recombinant channels. Both mutant MCU channels expressed at somewhat higher levels than that of rescue wild-type MCU by Western blot analyses of isolated mitochondrial lysates, with MICU1/2 dimer expression similar among the different cell lines (*SI Appendix Fig. S*2). However, with 1.5 mM EGTA and no added Ca^2+^ in the pipette solutions, Ca^2+^-current densities were not different between the mutant- and wild-type-MCU expressing cells (Fig. 1*A* and *B*), suggesting the functional expression of the mutant and wild-type channels was equivalent among the lines. Importantly, normal channel inhibition by 400 nM [Ca^2+^]_m_ (Fig. 1*A* and *B*) was abolished in both MCU-mutant channels (Fig. 3*A-C*). Furthermore, channel inhibition observed in high-[BAPTA]/low-[Ca^2+^]_m_ (Fig. 2*A* and *B*) was also abolished (Fig. 3*C*). In contrast, charge neutralization of E117, located in a distinct acidic patch in the amino terminus (28), in E117Q-MCU channels, was without effect on normal matrix [Ca^2+^] inhibition (*SI Appendix Fig. S*3*A*). These results indicate that the amino terminus of MCU contains a functional Ca^2+^-binding site encompassing residues D131 and D147 that accounts for matrix Ca^2+^ (and Mg^2+^) inhibition of channel activity. Occupancy of this site overrides normal cytoplasmic Ca^2+^-mediated MICU1/2 activation of channel activity to inhibit MCU. To independently test this, we measured intact-mitochondrial Ca^2+^ uptake in digitonin-permeabilized cells in response to incremental boluses of 5-7 µM Ca^2+^. Gatekeeping at low bath-[Ca^2+^] remained fully-functional in D131A-D147A- and E117Q-MCU-expressing cells (Fig. 3*E* and *SI Appendix Fig. S*4*A*), indicating that the MCU/EMRE/MICU1/2 channel complex remained intact and functional in the cells expressing mutant MCU. The rates of Ca^2+^ uptake in response to the first Ca^2+^ bolus were not different among the different cell lines (*SI Appendix Fig. S*4*B*), consistent with the electrophysiological data that indicated that the functional expression of wild-type and mutant MCU channels was equivalent. Whereas mitochondria in MCU-knockout cells rescued with wild-type MCU rapidly took up Ca^2+^ in response to 9-12 successive boluses, those in cells rescued with either D131A-MCU or D147A-MCU exhibited partial failure to take up Ca^2+^ after only two Ca^2+^ pulses, with complete failure observed after 3-4 pulses associated with collapse of ΔΨ_m_ (Fig. 3*E*), indicating matrix Ca^2+^ activation of the PTP. In contrast, cells rescued with E117Q MCU behaved similarly to wild-type MCU rescue cells (*SI Appendix Fig. S*3*B*). These results indicate that matrix Ca^2+^ regulation of MCU-channel activity observed in mitoplast electrophysiology is functional in intact mitochondria, with elevated matrix [Ca^2+^] feeding back to inhibit Ca^2+^ influx, mediated by the Ca^2+^-binding D131/D147 site in MCU. This feedback inhibition overrides cytoplasmic Ca^2+^-dependent MICU1/2 activation of MCU to prevent excessive mitochondrial Ca^2+^ uptake and ensure desensitization to deleterious PTP opening.

**Fig. 3.**
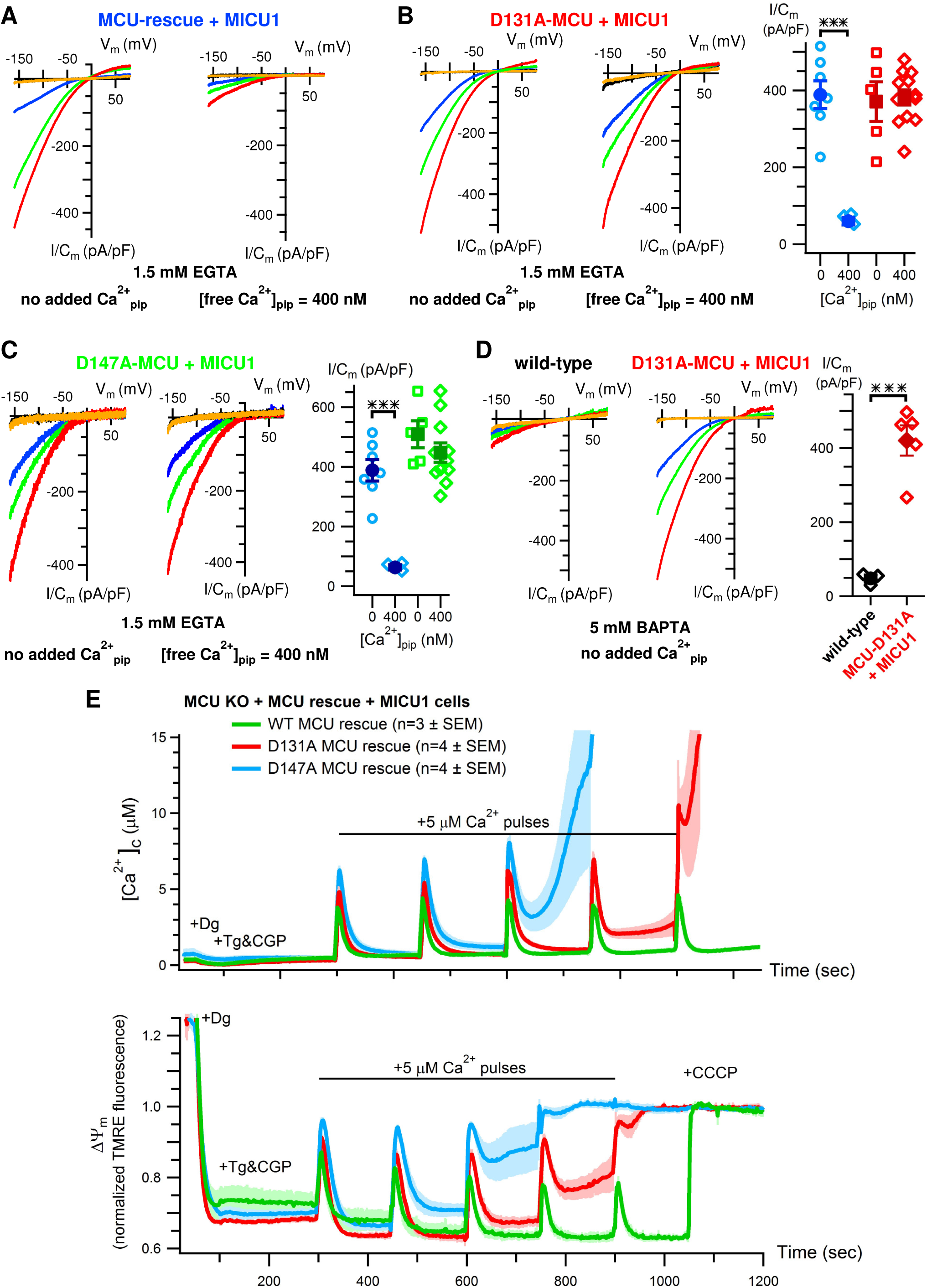
The MCU amino-terminus mediates matrix [Ca^2+^] inhibition of MCU. (*A*) Representative MCU Ca^2+^-current traces recorded from MCU-KO cells rescued with wild-type MCU and MICU1 with pipette buffer and [Ca^2+^] as indicated. (*B*) Left: similar in MCU-KO cells rescued with D131A-MCU (red label). Right: summary of MCU Ca^2+^ currents recorded in wild-type MCU and D131A-MCU mitoplasts with pipette solutions containing 0- or 400 nM free Ca^2+^ buffered with 1.5 mM EGTA. Bars, standard error of the mean (SEM); ***, *P* = 0.0005. (*C*), Similar, in MCU-KO cells rescued with D147A-MCU. Bars: SEM; ***, *P* = 0.0005. (*D*) MCU channel inhibition in 0-Ca^2+^/5 mM BAPTA (left; black symbols in summary panel at right; see also Fig. 2*A* and *D*) is abolished in mitoplasts expressing D131A-MCU (middle; red symbols in summary panel). Bars: SEM; ***, *P* = 0.0005. (*E*) Mitochondrial Ca^2+^ uptake and ΔΨ_m_ in digitonin-permeabilized cells in response to successive additions of 5-7 µM Ca^2+^ boluses in MCU-KO cells rescued with wild-type (WT) MCU and MICU1 (green; n=3) or with D131A- (red; n=4)) or D147A- (blue; n =4) MCU and MICU1. Traces show mean (solid line) ± SEM (shaded). Mitochondria in WT MCU-rescue cells are able to rapidly take up Ca^2+^ in response to 10-12 boluses (only 5 shown for comparison), whereas mutant MCU-rescue cells fail after 3-4 boluses, associated with collapse of ΔΨ_m_, indicative of matrix Ca^2+^-induced PTP opening.

### Regulation of MCU Na^+^ Currents

Whereas apo-MICU1/2 normally inhibits MCU-mediated Ca^2+^ influx in low [Ca^2+^]_i_ (13, 17-21), it fails to inhibit MCU Na^+^ currents observed in the absence of Ca^2+^ as the permeant ion (1, 22, 23). Furthermore, Na^+^ currents observed with matrix [Ca^2+^] buffered with 1.5 mM EGTA are independent of MICU1/2 association with the channel complex (23). We considered that Na^+^ currents are observed because they are recorded with pipette solutions lacking Mg^2+^ (1, 22, 23). However, addition of 2 mM Mg^2+^ to the pipette solutions did not inhibit the inward Na^+^ currents (Fig. 4*A*). Furthermore, whereas 300-400 nM Ca^2+^_m_ buffered with 1.5 mM EGTA inhibited Ca^2+^ currents (Fig. 1*A* and *B*), it was without effect on Na^+^ currents (Fig. 4*A*). In both conditions, the inhibitory D131/D147 Ca^2+^-sensor site was liganded, but it was no longer inhibitory with Na^+^ as the current carrier. Thus, under conditions in which Na^+^ is the current carrier, matrix [Ca^2+^]- and MICU1/2-regulation of MCU gating appear to be uncoupled. To explore this further, we examined the effects of high-[BAPTA]/low-[Ca^2+^]_m_ /0-Mg^2+^ pipette solutions. Notably, Na^+^ currents were strongly inhibited in this condition (Fig. 4*B*). Thus, with sufficiently strong Ca^2+^ buffering to ensure that all matrix Ca^2+^ sensors are unoccupied, normal apo-MICU1/2 inhibition of MCU activity is similarly observed for Ca^2+^ influx and Na^+^ currents.

**Fig. 4.**
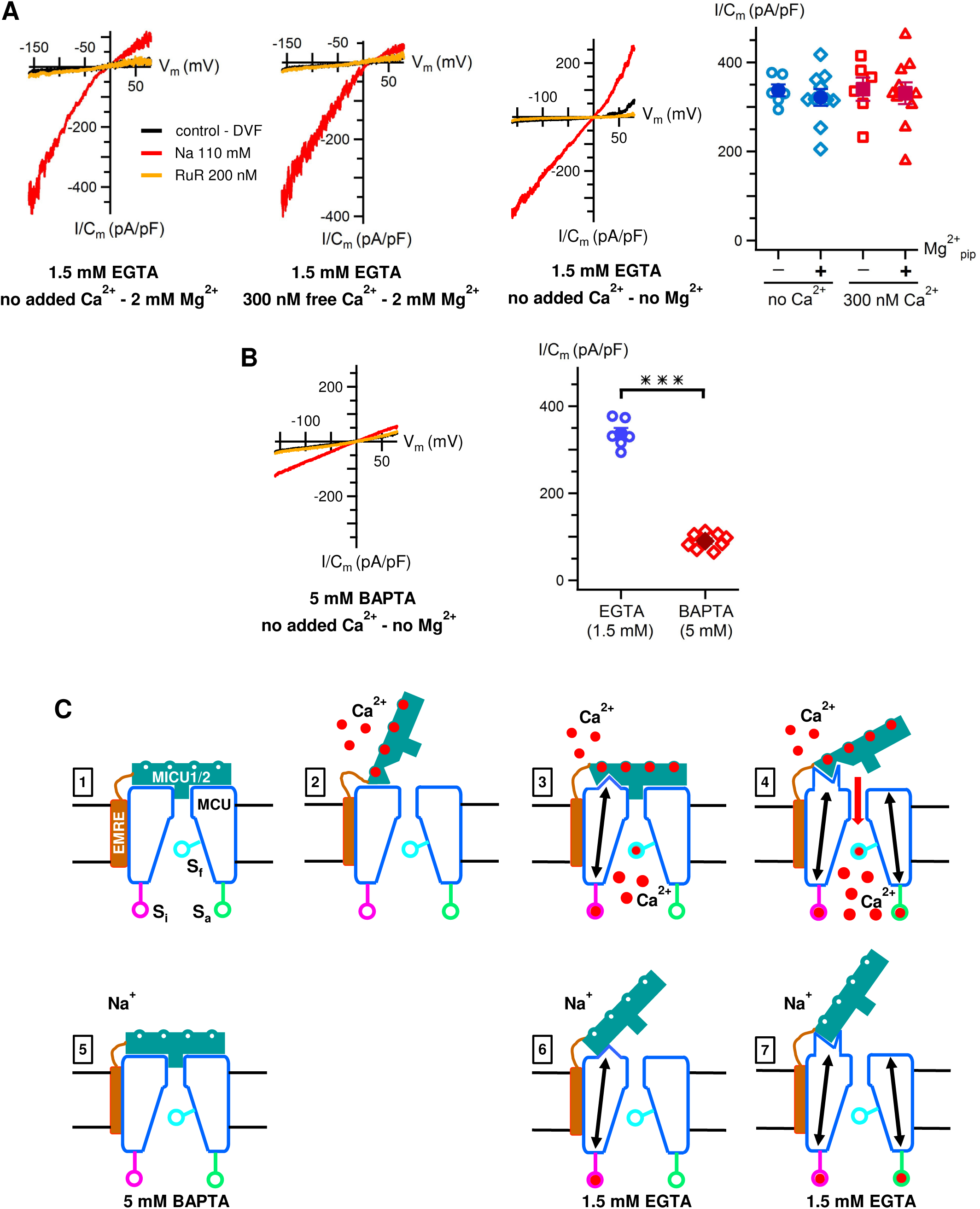
Regulation of MCU Na^+^ currents and a model for coupled matrix- and cytoplasmic-Ca^2+^ regulation. (*A*) MCU Na^+^ currents are not inhibited by matrix Ca^2+^ or Mg^2+^. First two panels: representative recordings of MCU-mediated Na^+^ currents in absence of added pipette Ca^2+^ (left) and presence of 300 nM matrix Ca^2+^ (right). Normal Ca^2+^-current inhibition by 300 nM matrix Ca^2+^ buffered with 1.5 mM EGTA (Fig. 1*A* and *B*) is not observed for MCU Na^+^ currents. Inward rectification of Na+ currents caused by Mg^2+^ in the pipette solution. Third panel: representative recordings of MCU-mediated Na^+^ currents in absence of both matrix Ca^2+^ (1.5 mM EGTA) and Mg^2+^. Lack of Mg^2+^ results in linear Na^+^ currents. Fourth panel: summary data with pipette solutions containing 1.5 mM EGTA with 0- (blue) or 300-400 nM (red) Ca^2+^ with and without 2 mM Mg^2+^. Data for 0-Mg^2+^/400 nM Ca^2+^ from reference (23). Bars: standard error of the mean (SEM). No significant differences among groups. (*B*) Left: representative Na^+^ current recordings with pipette solution containing 0-Mg^2+^ or 0-Ca^2+^/5BAPTA. Right: summary demonstrating inhibition of MCU Na^+^ currents by strong Ca^2+^ buffering in absence of matrix Mg^2+^. Bars: SEM; ***, *P* = 4 E-10. (*C*) Cartoon models of MCU-channel regulation by coupled MICU1/2 and matrix [Ca^2+^] mechanisms. Panels 1-4 depict physiological cytoplasmic- and matrix-[Ca^2+^] regulation of MCU-channel open probability. Panels 5-7 depict regulation of experimental MCU-mediated Na^+^ currents. MICU1 and MICU2 are depicted as a single entity (MICU1/2) facing the IMS with 4 Ca^2+^-binding sites. Distinct functional conformations of MICU1/2 and MCU are depicted, with comparable conformations in Ca^2+^- and Na^+^-permeation experiments aligned vertically. Arrows within MCU denote conformational changes in MCU associated with Ca^2+^ binding to inhibitory (S_i_), activating (S_a_) and flux (S_f_) sensor sites. See text for details.

## Discussion

Previous studies of intact mitochondria in permeabilized cells (10, 11, 13, 17-21) and in mice (29-31) and humans (32-34) have suggested important roles for IMS-localized MICU1/2 in MCU regulation by cytoplasmic [Ca^2+^]. Our results indicate that cytoplasmic MICU1/2 regulation of MCU is dictated by Ca^2+^ sensors located in the matrix, at least one of which is an integral component of the channel pore-forming subunit. A model that can account for our data is schematized in Fig. 4*C*. In the low-[Ca^2+^]_i_ regime characteristic of resting conditions in the cytoplasm, MICU1/2 in its apo-conformation maintains MCU in a closed-channel conformation (Fig. 4*C*[1]). Maintenance of this inhibited channel conformation may also be facilitated by matrix Ca^2+^, since we observed relatively strong inhibition of MCU Ca^2+^ currents even at 100 nM, the physiological resting matrix [Ca^2+^]. In the presence of elevated cytoplasmic [Ca^2+^], physiologically or when recording Ca^2+^ currents in mitoplast electrophysiology, Ca^2+^ binding to MICU1/2 induces a conformational switch (35-38) that relieves channel inhibition imposed by the apo-conformation, and mitochondrial Ca^2+^ uptake (10-13, 17-21) and Ca^2+^ currents (1, 22, 23, 39) can be recorded (Fig. 4*C*[2]). Ca^2+^ influx promotes Ca^2+^ binding to a flux sensor, which in turn tunes the affinities of other, coupled inhibitory and activating Ca^2+^-sensor sites in the matrix. The evidence for this was revealed by the biphasic matrix-[Ca^2+^] dependence of MCU Ca^2+^ current densities, and the effects of the fast Ca^2+^ buffer BAPTA in the matrix on MCU open probability. The biphasic nature of the relationship remained intact as BAPTA concentration was increased, suggesting that the inhibitory and activating sites were coupled and jointly regulated by a sensor that was sensitive to the matrix Ca^2+^ buffering capacity. Interestingly, divalent cation-flux feedback inhibition of CorA, the prokaryotic homolog of the primary mitochondrial Mg^2+^ channel Mrs2, is also regulated by dual divalent cation-binding sites in the channel cytoplasmic domain (40, 41). Because BAPTA at the same concentration as the slower Ca^2+^ buffer EGTA also shifted the matrix [Ca^2+^] dependence in a manner similar to that of simply increasing buffer capacity, we assume that the sensor that regulates the coupled inhibitory and activating sites is located within a region of high Ca^2+^ concentration that cannot be well buffered by EGTA. The source of this Ca^2+^ is most likely the Ca^2+^ that permeates through the channel pore. Accordingly, we have termed this the flux sensor. Its physical location remains to be determined, but we speculate that it could be an internal Ca^2+^-binding site associated within the channel selectivity filter (42) or localized at a putative inner gate at the bottom of the cavity below the pore (27). With the flux sensor and the inhibitory site fully Ca^2+^-liganded, the MCU channel open probability is strongly reduced even with MICU1/2 fully Ca^2+^-liganded (Fig. 4*C*[3]). In the presence of 1.5 mM EGTA, maximal inhibition of ∼80% is observed when matrix [Ca^2+^] is ∼400 nM. Two lines of evidence suggest that these observations in mitoplasts with artificial buffers are physiologically relevant. First, we were able to demonstrate that changes in the matrix [P_i_], the major mitochondrial matrix Ca^2+^ buffer *in situ*, also affected matrix Ca^2+^ regulation of MCU channel open probability, although these experiments were complicated by the low Ca^2+^ affinity of P_i_. Second, we demonstrated the existence and importance of matrix [Ca^2+^] regulation of MCU open probability *in situ* in intact mitochondria by Ca^2+^-uptake assays in permeabilized cells expressing mutant MCU channels that lacked matrix Ca^2+^ inhibition of mitoplast Ca^2+^ currents. These mitochondria underwent Ca^2+^ overload-induced PTP opening associated with a collapse of the ΔΨ_m_ in response to Ca^2+^ transients that were without detrimental effects on mitochondria expressing wild-type MCU with matrix Ca^2+^ inhibition intact. Importantly, inhibition of MCU channel open probability by matrix Ca^2+^ depends upon MICU1/2. The evidence for this was the lack of matrix [Ca^2+^] inhibition of MCU Ca^2+^ currents in cells lacking MICU1 or MICU2, and also in cells in which the heterodimer MICU1/2 was not functional due to the absence of its EMRE-mediated tethering to the channel complex. This suggests that Ca^2+^ binding to the matrix inhibitory site in the amino-terminus of MCU results in a conformational change in MCU that enables Ca^2+^-bound MICU1/2 to inhibit it (Fig. 4*C*[3]). In contrast, the data suggest that matrix Ca^2+^ binding to the lower-affinity activating site induces a different channel conformation that no longer accommodates Ca^2+^-liganded MICU1/2, enabling Ca^2+^ permeation (Fig. 4*C*[4]). Failure to account for the fundamental modulation by matrix [Ca^2+^] of MICU1/2 regulation may result in an inability to observe MICU1/2 regulation of Ca^2+^ currents by patch-clamp electrophysiology if the pipette [Ca^2+^] is either too high or too low, depending upon the matrix Ca^2+^-buffering properties. MICU1/2 regulation of Na^+^ currents was only revealed under conditions that strongly minimized occupancy of all matrix Ca^2+^ sensors (Fig. 4*C*[5]). Because this has not been previously considered (Fig. 4*C*[6,7]), it can explain why the channel has been observed to be constitutively active when Na^+^ currents are measured (1, 22, 23).

Our results indicate that Ca^2+^ occupancy of matrix Ca^2+^ sensors regulates MCU channel-conformational states that determine the efficacy of MICU1/2 regulation. Previously, regulation of MCU channel activity was defined by cytoplasmic Ca^2+^ modulation of MICU1/2 that sets a [Ca^2+^]_i_ threshold for MCU channel activation and its cooperative activation. Our results indicate that this regulation is set not only by the Ca^2+^ affinities of MCU1/2 EF hands, but also by matrix [Ca^2+^] and buffering capacity, allowing for enhanced cellular regulation of mitochondrial Ca^2+^ homeostasis. Recent studies have demonstrated a direct biochemical interaction of MICU1/2 with the MCU selectivity filter (15, 16). Because mutations in either MCU or MICU1 that disrupt this interaction lead to a constitutively open channel, it has been suggested that MICU1/2 functions as a classic pore blocker (16). Our results here indicate that structural changes that affect this interaction are localized not only in MICU1/2 (35-38), regulated by cytoplasmic Ca^2+^, but also in the MCU channel itself, regulated by Ca^2+^ sensors localized on the opposite side of the IMM. This mechanism is reminiscent of Ca^2+^-dependent inactivation of plasma membrane voltage-gated Ca^2+^ channels (VGCC). There, Ca^2+^ flux through the channel pore interacts with channel elements in the cytoplasm that result in structural changes in the selectivity filter on the other side of the membrane that close the channel (43). Similarly, voltage-dependent inactivation of VGCC involves elements in the cytoplasmic aspect of the channel that drive conformational changes close to the ion-conducting pore (43, 44). Alternately, it has been suggested that MCU may possess an internal gate at the base of the internal central cavity below the selectivity filter (27). It is possible that structural changes regulated by MICU1/2 and matrix Ca^2+^ sensors regulate this gate. An important goal now is to understand how Ca^2+^ binding to mitochondrial matrix sensors are transduced into conformational changes of the MCU channel pore region or internal gate that affect MCU interactions with MICU1/2 to regulate Ca^2+^ permeation.

## Materials and Methods

### Cell lines, Constructs and Antibodies

Wild-type HEK-293T and KO cell lines were grown in Dulbecco’s modified Eagle’s medium (DMEM) supplemented with 10% FBS, 100 U/ml penicillin, and 100 µg/ml streptomycin at 37 °C and 5% CO_2_. MCU-KO, MICU1-KO and MICU2-KO HEK-293T cells were a generous gift from Dr. Vamsi Mootha. Full length MICU1, MICU2 and EMRE cDNAs were purchased from Origene Technologies USA. EMRE-ΔC-V5 constructs were generated by a PCR-based cloning strategy as described (23). All constructs were sequence-verified and stably transfected into KO backgrounds. D131A-MCU-Flag and D147A-MCU-Flag cDNAs in pCMV6-A-BSD were a gift from Dr. Muniswamy Madesh. E117Q-MCU-V5-His in pCMV6-A-Puro was generated using the QuickChange II XL site-directed mutagenesis kit and XL10-Gold® ultracompetent *E. coli*. All MCU constructs were sequence-verified and stably transfected in the MCU-KO backgrounds together with MICU1. MCU WT-rescue or mutant cells were grown in complete DMEM supplemented either with blasticidin (5 µg/ml), puromycin (2 µg/ml), or both, until stable expression of each plasmid was achieved. Clones were isolated by limiting dilution for each condition and maintained in complete medium with antibiotics. Exclusive mitochondrial localization was verified by immunofluorescence. Only clones expressing recombinant proteins in all mitochondria in all cells were selected for experimentation.

### Uniporter Electrophysiology

Mitoplast electrophysiology was performed as described (1, 22, 23). Patch pipettes had resistances of 20 - 60 MΩ when filled with (in mM): 120 TMA-OH, 120 HEPES, 80 D-gluconic acid, 10 glutathione; 2 MgCl_2_; pH = 7.0 with D-gluconic acid; osmolarity 390-450 mOsm/kg. EGTA or BAPTA were used, as specified, to buffer correspondent amounts of CaCl_2_ for [Ca^2+^]_free_ ≤ 400 nM (confirmed by Ca^2+^-sensitive dye fluorimetry). Mitoplasts were initially bathed in (in mM): 150 KCl, 10 HEPES, 1 EGTA; pH = 7.2; osmolarity 300 mOsm/Kg (“KCl-DVF” solution). Voltage pulses (350 - 500 mV for 15 - 50 ms), were delivered by the PClamp-10 (Molecular Devices) program to obtain the “whole-mitoplast” configuration. Access resistance (30 - 90 MΩ) and mitoplast capacitance C_m_ (0.2 - 1 pF) were determined using the membrane test protocol of the PClamp-10 software. After the whole-mitoplast configuration was obtained, the KCl-DVF bath solution was exchanged with: HEPES-EGTA [(in mM): 150 HEPES, 1.5 EGTA; pH = 7.0 with Tris-base] for baseline measurements, and then replaced with HEPES-0-EGTA solutions with 0.1, 0.3, or 1 mM CaCl_2_ successively. Finally, a HEPES-based solution with 1 mM CaCl_2_ and 200 nM ruthenium red (RuR) was perfused into the bath to record the final baseline (= I_RuR_) after block of the MCU currents. Osmolarities of bath solutions were 297-305 mOsm/Kg, adjusted with sucrose. The voltage protocol, delivered by the PClamp-10 software with a DigiData-1550 interface (Molecular Devices), consisted in stepping from V_m_ = 0 mV to -160 mV for 20 ms, followed by ramping to 80 mV (rate of 279 mV/s), dwelling at 80 mV for 20 ms, and returning to 0 mV. Currents were recorded using an Axopatch 200-B amplifier at room temperature with a sampling rate of 50 kHz and anti-aliasing filtered at 1 kHz. Data analysis was performed with the PClamp-10 software. For quantitative comparisons, current densities are defined as: *I*_*MCU*_ */ C*_*m*_ *= (I*_*Ca*_ *– I*_*RuR*_*) / C*_*m*_, with *I*_*Ca*_ and *I*_*RuR*_ measured at *V*_*m*_ = -160 mV in the presence of 1 mM Ca^2+^ in the bath.

### Simultaneous Measurements of Mitochondrial Ca^2+^ Uptake and Mitochondrial Membrane Potential in Permeabilized Cells

Concurrent measurements of mitochondrial Ca^2+^ uptake and IMM potential (ΔΨ_m_) in permeabilized HEK-293T cells were performed as described (18). Cells were grown in 10-cm tissue culture coated dishes for 48 hr prior to each experiment. 6-8×10^6^ cells were trypsinized, counted, and washed in a Ca^2+^-free extracellular-like medium, centrifuged, suspended in 1.5 ml of intracellular-like medium (ICM - in mM: 120 KCl, 10 NaCl, 1 KH_2_PO_4_, 20 HEPES, and 5 succinate [pH 7.2]) that had been treated with BT Chelex^®^ 100 resin before use to attain an initial free [Ca^2+^] of ∼20 nM, and transferred to a cuvette. The cuvette was placed in a temperature-controlled (37°C) experimental compartment of a multi-wavelength-excitation dual wavelength-emission high-speed spectrofluorometer (Delta RAM, Photon Technology International). Membrane-impermeable Fura2 K^+^ salt (K_D_ = 140 nM) or FuraFF K^+^ salt (K_D_ = 5.5 µM) (final concentration 1 µM) and TMRE (final concentration 1 µM) were added at t=25 sec from the start, to measure bath [Ca^2+^]_c_ and ΔΨ_m_ concurrently. Fluorescence at 549-nm excitation/595-nm emission for TMRE, along with 340-nm and 380-nm excitation/535-nm emission for Fura2/FuraFF, was measured at 5 Hz. 40 µg/mL digitonin was added at t=50 sec to permeabilize cells and allow the cytoplasm to equilibrate with the bath solution such that the degree of quenching of TMRE fluorescence reports the relative ΔΨ_m_. Fura2/FuraFF fluorescence under these conditions is related to the cytoplasmic [Ca^2+^] ([Ca^2+^]_c_) according to 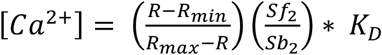, where (*R*) is the measured fluorescence ratio (340/380 nm), (*R*_*min*_) is the fluorescence ratio under 0-[Ca^2+^], (*R*_*max*_) is the fluorescence ratio under saturating [Ca^2+^], (*Sf*_*2*_) and (*Sb*_*2*_) are the absolute 380 nm excitation/535 nm emission fluorescence under 0- [Ca^2+^] and saturating [Ca^2+^], respectively, and (*K*_*D*_) is the dissociation constant of the dye for Ca^2+^. Thapsigargin (2 µM) and CGP37157 (20 µM) were added at 100 sec to inhibit Ca^2+^ uptake into the endoplasmic reticulum and NCLX-mediated mitochondrial Ca^2+^ extrusion, respectively. After [Ca^2+^]_c_ reached a steady state at ∼300 sec, MCU-mediated Ca^2+^ uptake was initiated by adding a bolus of 5 µl of 5 mM CaCl_2_ to the cuvette to achieve increases in [Ca^2+^]_c_ between ∼5 µM. The volume of CaCl_2_ required to achieve the desired increase in [Ca^2+^]_c_ was calculated based on the activity coefficient for Ca^2+^ in ICM. After [Ca^2+^]_c_ was monitored for 150 sec following Ca^2+^ addition, a second bolus of CaCl_2_ was added. This procedure was repeated for a total of five to nine successive additions of Ca^2+^. 150 sec after addition of the final bolus, CCCP (2 µM) was added to uncouple ΔΨ_m_ and allow unimpeded Ca^2+^ efflux from mitochondria as a measure of the total extent of uptake. Under our conditions, each Ca^2+^ bolus caused a transient depolarization as measured with TMRE that recovered after ∼50-60 sec. Concurrent failure of Ca^2+^ uptake and loss of ΔΨ_m_ indicates opening of the mitochondrial permeability transition pore (PTP).

Initial Ca^2+^ uptake rates in permeabilized cells were determined for each experiment using single-exponential fits from the time of the first Ca^2+^ addition (t = 0 sec) to achievement of a new steady state (t = 300 sec) to obtain parameters **A** (extent of uptake) and **τ** (time constant) (IGOR Pro; RRID:SCR_000325). The instantaneous rate of uptake (R) at t = 0 is equal to the first derivative of the fit, and standard deviations (SD) of rates were calculated from standard deviations of A (SD_A_) and τ (SD_τ_) as follows:

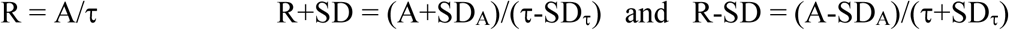

For determination of the steady-state [Ca^2+^]_c_ after completion of uptake (gatekeeping threshold [Ca^2+^]_i_), the calculated [Ca^2+^]_i_ 300 sec after challenge with 0.1 - 10 µM Ca^2+^ (**Y**) was plotted as a function of the (peak) initial [Ca^2+^]_i_ achieved after bolus Ca^2+^ addition (**X**). Data were fitted using a one-phase association model where (**Y**_**0**_) is the Y value when X is zero, (**P**) is the plateau, or Y value at infinite X, and (**K**) is the rate constant, expressed in reciprocal of the X axis.

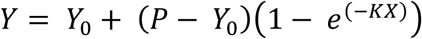

The value of (**P**) represents to the steady-state [Ca^2+^]_i_ at which gatekeeping is re-established +/- the standard error of (**P**) determined from the fit.

### SDS-PAGE western blotting

For western blots of mitochondrial proteins, mitochondria were isolated as described (6). Total protein concentrations were calculated using BCA Protein Assay kit (Thermo Fisher Scientific Cat# 23225), and samples for PAGE were prepared in 2x Laemmli sample buffer (Bio-Rad Cat# 1610737) +/- 2-Mercaptoethanol (βME) (Bio-Rad Cat# 1610710). NuPAGE© gels (4-12% - Thermo Fisher Scientific Cat# NP0321) were transferred to Immobilon© P PVDF membranes (Millipore-Sigma Cat# IPVH00010) and probed with various antibodies. Antibodies used: anti-MCU (Cell Signaling Technology Cat# 14997S), anti-V5 tag (Cell Signaling Technology Cat# 13202S), anti-MICU2 (Abcam Cat# ab101465), anti-DYKDDDDK (Flag tag) (Cell Signaling Technology Cat# 14793S), anti-MICU1 (Cell Signaling Technology Cat# 12524S) anti-β-Tubulin (Invitrogen Cat# 32-2600), and anti-Hsp60 (Abcam Cat# ab46798). Membranes were blocked in 5% fat-free milk for 1 hr at RT, incubated overnight at 4°C with primary antibody, and then for 1 hr at RT with anti-rabbit IgG-HRP (Cell Signaling Technology Cat# 7074S) or anti-mouse IgG-HRP (Cell Signaling Technology Cat# 7076S) secondary antibodies conjugated to horseradish peroxidase (HRP). Chemiluminescence detection was carried out using SuperSignal West Chemiluminescent Substrate (Thermo Fisher Scientific Cat# 34580). The mean pixel density of bands was quantified using the Fiji software package in ImageJ (ImageJ, RRID: SCR_003070), corrected for the intensity of the loading-control band (β- tubulin for whole-cell lysates or Hsp60 for isolated mitochondria), and normalized to the response in WT cells.

## ACKNOWLEGEMENTS

We thank Drs. Francesca Fieni and Yuri Kirichok for advice regarding mitoplast electrophysiology, Usha Patel for assistance with molecular biology, and Dr. Don-On Daniel Mak for helpful discussions. Supported by NIH R37 GM56328.

## Author contributions

H.V. performed the electrophysiology. R.P. performed molecular biology, biochemistry and creation of stable cell lines. R.P. and C.L. performed functional studies of permeabilized cells. J.K.F. supervised the project. H.V. and J.K.F wrote the manuscript.

## Additional information

**SI Appendix** consists of 4 figures.

**Correspondence and requests for material** should be addressed to J.K.F.

