## Supplemental Figures for "Strongly coupled transmembrane mechanisms control MCU-mediated mitochondrial Ca^2+^ uptake"

### Supplementary Appendix Figure Legends

**SI Appendix Fig. 1.** Matrix phosphate ($P_i$) modulates MCU activity. (*A*) Cartoon depicting rationale for the experimental design to examine the effects of $P_i$ on matrix $Ca^{2+}$ regulation of MCU $Ca^{2+}$ channel activity imposed by poor $Ca^{2+}$ buffering properties of $P_i$ ($K_d$ ~ 70 μM). Assuming only that the biphasic response of MCU activity to $[Ca^{2+}]_m$ shifts left with increasing $[P_i]$, a change in MCU activity could become apparent if the working $[Ca^{2+}]$ was chosen to lie either on the descending or ascending phase, but not on the plateau (both possibilities indicated by dashed lines). (*B*) MCU $Ca^{2+}$ currents were recorded with pipette solution containing 2 μM free $Ca^{2+}$ (indicated at top) with various $[P_i]$ given under each panel. (*C*) MCU $Ca^{2+}$ currents were recorded with pipette solution containing 500 nM free $Ca^{2+}$ with various $[P_i]$. MCU activity with $[P_i]_{pip}$ = 1 μM was reduced relative to that in 3 or 10 μM $[P_i]_{pip}$ (***, $P = 0.002$). With $[P_i]_{pip}$ = 1 μM and 790 nM $[Ca^{2+}]_{pip}$, MCU activity was maximal, suggesting that data at 500 nM $Ca^{2+}$ corresponded to the ascending phase of the biphasic MCU dependence on matrix $[Ca^{2+}]$. Bars: standard error of the mean; **, $P = 0.008$.

**SI Appendix Fig. 2.** Expression of MCU and MICU1/2 in MCU-rescue cells. Western blot analyses of mitochondrial lysates from stable HEK293 cell clones derived from MCU-KO cells rescued with wild-type MCU and MICU1, or mutant E117Q-, D131A- or D147A-MCU and MICU1. MCU (left) and MICU1-MICU2 heterodimer (right) levels and Hsp60 as loading control shown. Quantification of band intensities, normalized to loading control and to wild-type MCU levels, shown at bottom. Bars: standard error of the mean; n = 3 separate mitochondrial preparations from each cell line.

**SI Appendix Fig. 3.** E117Q-MCU behaves similarly to wild-type MCU. (*A*) Left: representative MCU $Ca^{2+}$ currents recorded from E117Q-MCU mitoplasts with 1.5 mM EGTA and either no-added $Ca^{2+}$ or 400 nM free $Ca^{2+}$ in the pipette solution. Right: summary of MCU $Ca^{2+}$-current densities in mitoplasts from MCU-KO cells rescued with wild-type MCU and MICU1 (blue) or E117Q-MCU and MICU1 (green) with pipette $[Ca^{2+}]$ as indicated. Bars: standard error of the mean; ***, $P = 0.0001$. (*B*) Responses to 5 µM $Ca^{2+}$ boluses induced rapid mitochondrial $Ca^{2+}$ uptake similarly in wild-type MCU rescue and E117Q-MCU rescue cells. Dg: 40 µg/ml digitonin; Tg: 2 µM thapsigargin; CGP: 20 µM CGP3715.

**SI Appendix Fig. 4.** Mitochondrial $Ca^{2+}$ uptake in permeabilized MCU-KO cells stably rescued with wild-type and mutant MCU. (*A*) Gatekeeping threshold cytoplasmic $[Ca^{2+}]$ ($[Ca^{2+}]_c$) derived from one-phase association fits of steady-state $[Ca^{2+}]_c$ after addition for each cell line measured 300 sec after first bolus of 5 µM $Ca^{2+}$. (*B*) Initial $Ca^{2+}$ uptake rates in response to first 5 µM $Ca^{2+}$ bolus addition. Each point represents an independent experiment. Bars: standard error of mean. No differences among groups.

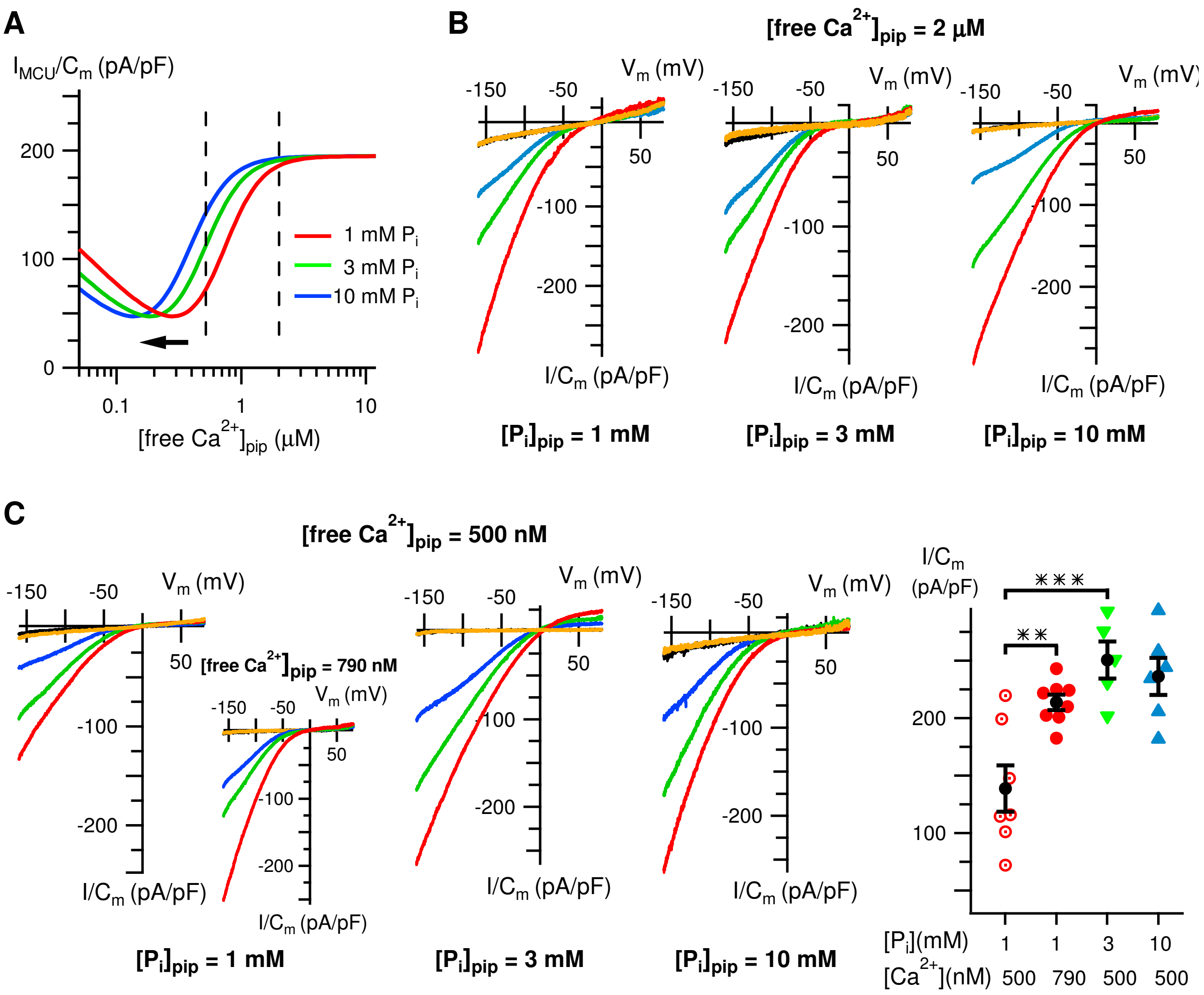


**SI Appendix figure 1**

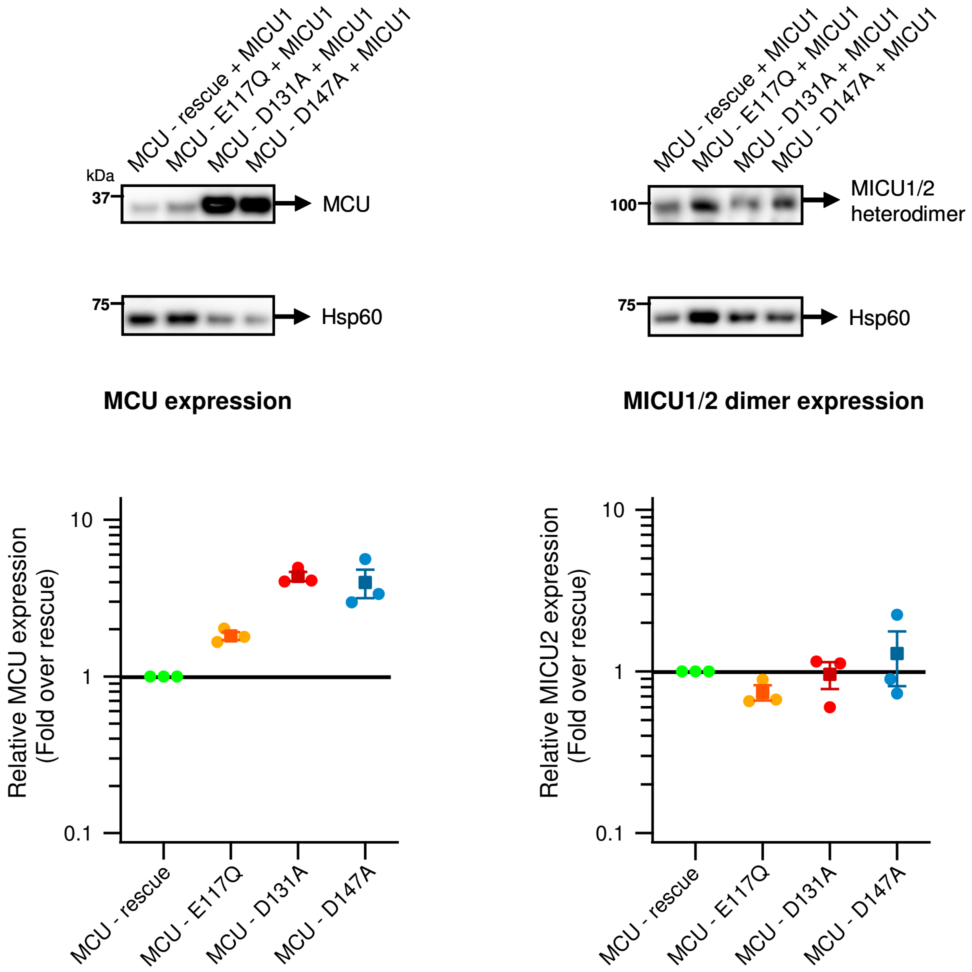


**SI Appendix Figure 2**

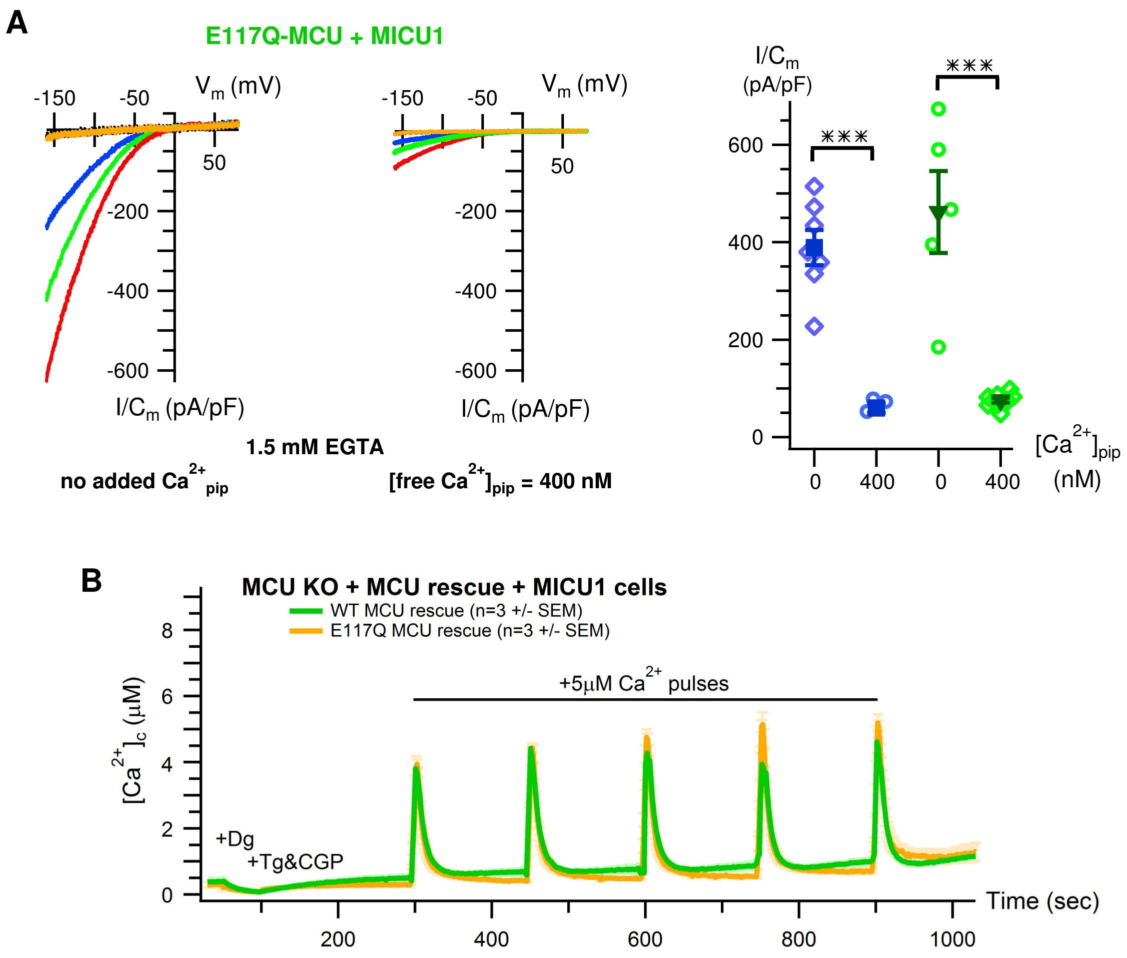


SI Appendix figure 3

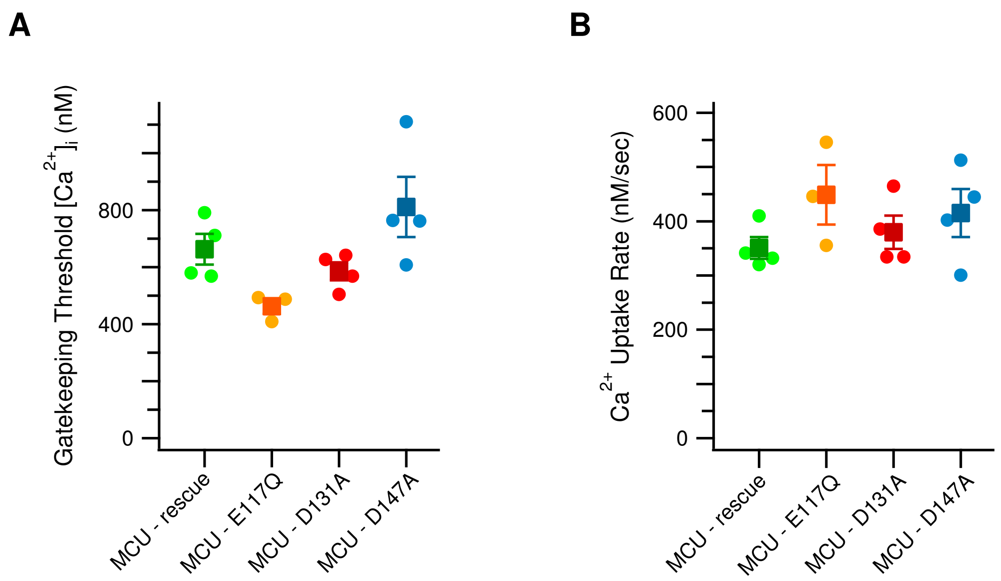


**SI Appendix figure 4**
